# CPP2Vec: a Representation Learning Approach for Cell-Penetrating Peptides Prediction

**DOI:** 10.1101/2025.05.15.654208

**Authors:** Stavroula Svolou, Vasileios Konstantakos, Anastasia Krithara, Georgios Paliouras

## Abstract

**Background:** Cell-penetrating peptides (CPPs) facilitate the delivery of a variety of therapeutic molecules across the plasma membrane, from small chemical substances to nucleic acid-based macromolecules, such as antisense oligonucleotides (ASOs). Among neutral ASOs, peptide nucleic acids (PNAs) and phosphorodiamidate morpholino oligomers (PMOs) have been extensively studied as potential medical treatments for Duchenne Muscular Dystrophy (DMD), a severe genetic disease that causes muscle degeneration progressively. Over the last few decades, many *in silico* methods have emerged to detect novel CPPs, counterbalancing the cost of wet-lab experiments.

**Results:** In this study, we propose CPP2Vec, a Word2Vec-based CPP prediction method, where the Word2Vec technique is used to represent amino acid sequences of peptides. We developed three task-specific supervised machine learning models for CPP-Classification, Uptake-Efficiency and PMO-Delivery. The first two models were designed to determine if an unseen peptide is a CPP and to predict its uptake efficiency, respectively, while the PMO-Delivery model predicts if a peptide could enhance the cellular delivery of a PMO-complex compared to its naked version. Furthermore, we explored an alternative approach using pre-trained protein-based Large Language Models (LLMs) - T5, BERT, and ESM-2 - to generate the embeddings, resulting in three task-specific models, namely CPP2LLM. A comparison of CPP2Vec and CPP2LLM with state-of-the-art CPP prediction tools is included, proving their significant predictive performance.

**Conclusion:** In this research, we present a Machine Learning (ML)-based tool that introduces the use of the Word2Vec technique in the field of CPPs prediction. Notably, it stands out for not requiring any manual *a priori* feature engineering and for its ability to generalize without any changes between studied tasks. CPP2Vec is available for use at: https://github.com/SSvolou/CPP2Vec.

## Introduction

As the therapeutic landscape expands at an increasing rate, the need to design drug delivery methods to improve therapeutic efficacy, minimize toxicity side effects, and enable innovative medical treatments emerges.

To this end, CPPs have been introduced to facilitate the delivery of a variety of therapeutic molecules across the plasma membrane, from small chemical substances to nucleic acid-based macromolecules. CPPs are short peptides comprised of 5-30 amino acids that are categorized based on their origin, conformation, and physical-chemical character into subgroups [1]. Even though the evolution of next-generation sequencing technologies accelerates peptide sequencing, the cost of wet-lab experimental studies remains high in terms of both time and resources.

For this reason, in the last few decades, many in silico - mainly ML-based - methods have been developed to detect novel CPPs, counterbalancing the cost of conventional in vitro assays [2]. Some of them, not only predict if a peptide is a CPP or not, but also categorize them depending on their predicted uptake efficiency into “high” or “low” classes. A crucial component in the development of prediction models is the technique that has been used for the sequence representation. Currently available tools use some conventional feature descriptors aiming to capture compositional or physicochemical properties of CPPs, such as amino acid composition (AAC), composition of K-spaced amino acid pairs (CKS) and pseudo amino acid composition (PseAAC) [3–6], while others are based on binary encoding [7].

A summary of the existing ML-based CPP prediction tools is provided in Table 1. Most of the approaches use Random Forest as the primary ML method, while half of them explicitly emphasize the classification of CPPs. Only four of them are capable of handling both CPP-Classification and Uptake-Efficiency tasks (i.e., CPPred-RF, MLCPP, Wolfe et al., and MLCPP 2.0), while so far only one was designed to predict PMO-Delivery efficacy.

**Table 1.** Brief overview of existing state-of-the-art ML-based prediction tools.

| Year | Name | Method | CPP Classification | Uptake Efficiency | PMO Delivery |
| --- | --- | --- | --- | --- | --- |
| 2013 | CellPPD [7] | SVM | ✓ | × | × |
| 2016 | Diener et al. [14] | RF - SVM | ✓ | × | × |
| 2017 | SkipCPP-Pred [4] | RF | ✓ | × | × |
| 2017 | CPPred-RF [5] | RF | ✓ | ✓ | × |
| 2018 | MLCPP [15] | Extra Trees - RF | ✓ | ✓ | × |
| 2018 | KELM-CPPpred [3] | Kernel-based Extreme ML | ✓ | × | × |
| 2018 | CPPred-FL [6] | RF | ✓ | × | × |
| 2018 | Wolfe et al. [16] | RF | ✓ | ✓ | ✓ |
| 2022 | MLCPP 2.0 [17] | Stacked Ensemble Learning | ✓ | ✓ | × |
| 2025 | **CPP2Vec** | RF - SVM | ✓ | ✓ | ✓ |

SVM: Support Vector Machine. RF: Random Forest Classifier. The tool described in this work is marked in bold.

In this study, we introduce the usage of LLMs to produce contextualized embeddings for amino acid sequences in the field of CPP prediction. We evaluate the performance of the pre-trained LLMs: T5 [8], BERT [9] and ESM-2 [10], while simultaneously we introduce CPP2Vec, a Word2Vec-based CPP prediction method, where the Word2Vec (W2V) [11] technique is used to generate the representations of amino acid sequences of peptides. Our proposed method not only classifies uncharacterized peptides into CPP/non-CPP and high/low uptake efficiency categories, but also provides a reliable model predicting if a peptide could enhance the delivery of a phosphorodiamidate morpholino oligomer (PMO)-complex into the cell compared to its naked version. Finally, we include a case study for DMD [12, 13] highlighting CPP2Vec’s potential usage in real-life scenarios.

## Material and Methods

In this section, we summarize the approach that we followed to construct our proposed prediction models. Specifically, we describe our 4-stage CPP prediction pipeline, providing details about the selected training and test datasets, as well as information about how we evaluated the performance of our method.

### Description of CPP prediction pipeline

The workflow we implemented for CPP prediction is illustrated in Figure 1 and includes four main stages. The first stage involves dataset selection and preparation. To construct a high-quality prediction model, reliable and rigorous training and test datasets were needed to ensure unbiased and accurate results. The second stage is the generation of contextualized embeddings for CPP candidates. In this study, we utilized two approaches to represent amino acid sequences in a vectorized format. First, we employed a dataset-specific W2V technique, and afterward, we utilized the pre-trained protein LLMs: T5 [8], BERT [9], and ESM-2 [10]. The third stage is ML model construction. Various ML algorithms, including Support Vector Machine (SVM), Random Forest (RF), and Gradient Boosting (GB) classifiers, were tested to detect the best model for each task. Finally, the fourth stage involves the evaluation of the proposed models to confirm that the predicted results are trustworthy. A comparison of our approach with state-of-the-art tools was also conducted during this stage.

**Fig. 1.**
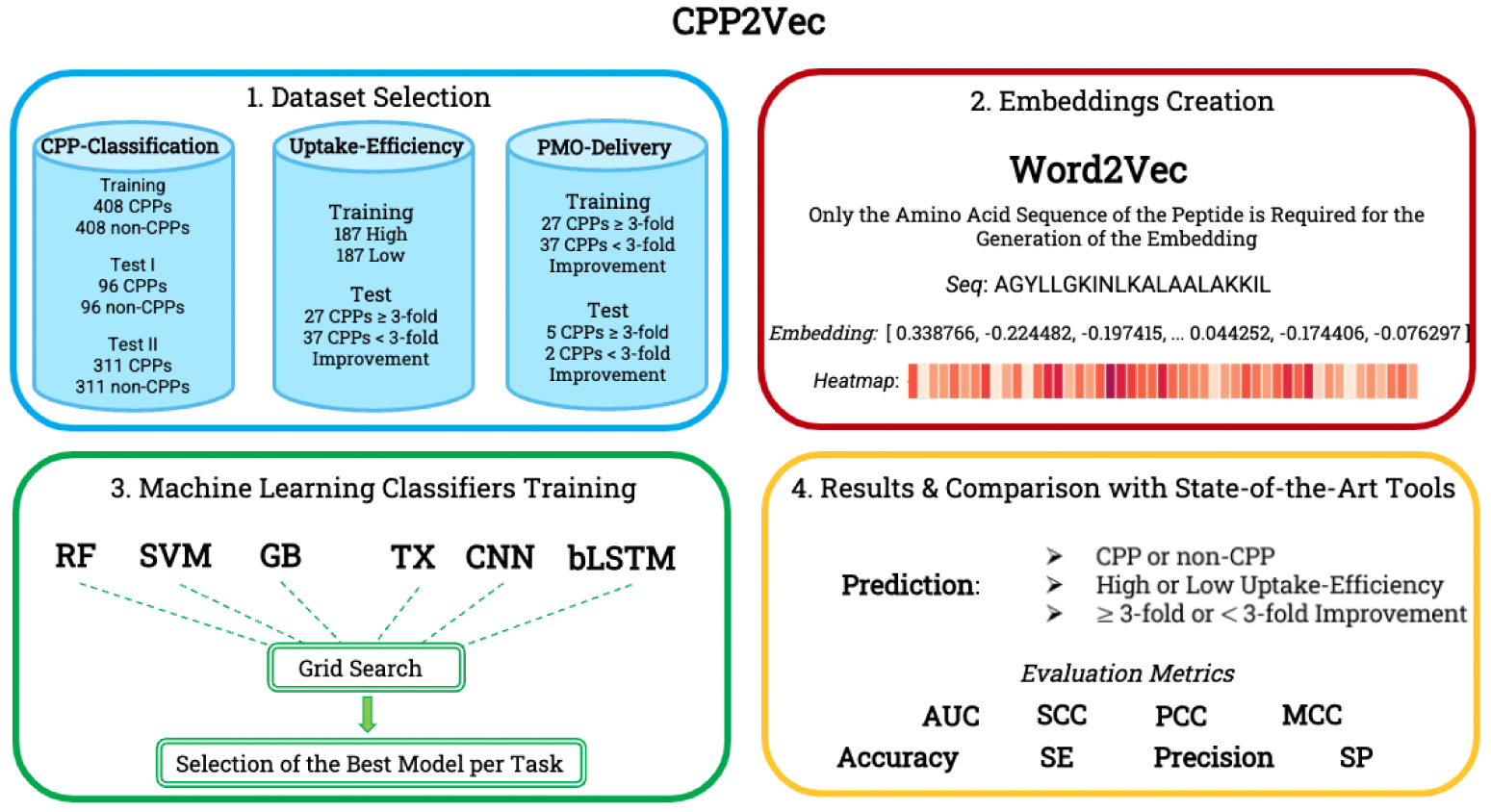
Workflow of CPP2Vec construction. (1) Datasets selection and data preprocessing. (2) Generation of contextualized embeddings through dataset-specific W2V approach. (3) Testing various ML algorithms to detect the proper one for each task. (4) Evaluation of the proposed models and comparison with state-of-the-art tools.

### Preparation of datasets

Considering each of the tasks that we studied (i.e. CPP-Classification, Uptake-Efficiency and PMO-Delivery), we carefully selected training and test datasets. For the sake of consistency, we needed to make an assumption to ensure fair comparisons with state-of-the-art tools. Since none of these tools accept non-natural amino acids in the provided sequences, we substituted *β*-alanine with *α*-alanine and 6-aminohexanoic acid with lysine in each of the involved datasets, respectively [18].

### Construction of training datasets

For the CPP-Classification task, we selected the KELM-CPPpred main dataset for training [3]. This dataset includes 408 non-redundant positive peptides derived from the CPPsite 2.0 database [19]. Negative samples were generated by combining 374 randomly selected non-redundant peptides from *CAMP_R_*_3_ [20] and BIOPEP [21] databases, along with 34 experimentally validated non-CPPs from the study of Sanders et al. [22] To reduce sequence redundancy, the cd-hit [23] was used with a threshold value of 0.8. The final dataset consists of 408 CPPs and an equal number of non-CPPs. For the Uptake-Efficiency task, we utilized for training the CPPsite3 dataset, pro-posed by Gautam et al. [7] in 2013. This dataset is derived from the CPPsite database [24], the predecessor of the CPPsite 2.0 database that contains 843 experimentally validated CPPs from which 430 were derived from research publications, 172 from patents and 241 from both of them. The majority of them (522) are protein derived while the rest of them are either synthetic (278) or chimeric (43). From these, peptides containing non-natural amino acids or D-amino acids were excluded due to their rare involvement in most biological systems, resulting in a dataset of 708 unique peptides. This step aims to maintain dataset’s uniformity and ensure that predictions will be more applicable to typical natural scenarios. Since these CPPs have been tested for their cell penetration capability under different cell types and conditions, authors classified them based on their uptake efficiency into three categories (i.e. low, medium and high), where high CPPs score *>*75% relative to control sample. Considering that only few peptides have been experimentally validated as non-CPPs, the “high” labelled peptides were combined with randomly generated peptides from SwissProt [25] to form the CPPsite3 dataset. Specifically, CPPsite3 contains 187 high uptake efficiency CPPs and an equal number of low-uptake efficiency CPPs (374 peptides in total).

For the PMO-Delivery task, our proposed model aims to determine whether conjugating a given peptide would enhance PMO activity by at least 3-fold. In order to accomplish this, we selected as our training set the 64-peptides dataset generated by Wolfe et al. [16] For our classification purposes, we assigned label 1 to peptides that demonstrated a 3-fold or greater improvement in eGFP fluorescence compared to unconjugated PMOs, while assigning label 0 to others, a concept initially described in the same study.

### Independent test datasets

For the CPP-Classification task, we validated the performance of our proposed model using two independent datasets: KELM-CPPpred [3] Independent and MLCPP [15] Independent datasets. For the sake of simplicity, they will be denoted as *kelm* and *mlcpp*, respectively.

The *kelm* dataset comprises 99 CPPs collected manually from the available bibliography, and 99 non-CPPs produced by SwissProt [25]. The sequences in this dataset are not included in the training dataset, while sequences showing *>*80% similarity were removed by the authors. The final dataset consists of 96 CPPs and an equal number of non-CPPs.

Concerning the *mlcpp* dataset, positive samples were extracted from CPPsite 2.0 [19], while negative samples were generated from SwissProt [25]. During the production of negative samples, peptides sharing similarities with known CPPs were excluded, and the remaining peptides were considered as non-CPPs. The cd-hit [23] program was also employed in this dataset to reduce sequence redundancy, using the same threshold value. The final dataset consists of 311 CPPs and 311 non-CPPs (622 peptides in total).

For the Uptake-Efficiency task, we evaluated our proposed model using Wolfe et al.’s [16] 64-peptides adjusted dataset with labels 1 and 0. The final dataset comprises 27 peptides with high uptake efficiency and 37 peptides with low uptake efficiency.

Finally, for the PMO-Delivery task, we utilized as a test set the 7 peptides proposed by Wolfe et al. [16] in the same study. These peptides were synthesized and experimentally tested in the fluorescence reporter assay. The dataset consists of 5 peptides with at least a 3-fold improvement and 2 with less than a 3-fold improvement.

### Generation of sequence embeddings

Driven by the parallels between amino acid sequences and natural languages, scientists have utilized various techniques to represent peptides in a vectorized format. In this study, we explore two approaches – the W2V method and pre-trained LLMs – in the realm of CPP prediction.

### Word2Vec approach

In recent years, the W2V technique [11] has been extensively utilized to generate word embeddings in the field of Natural Language Processing (NLP), demonstrating its robust performance. Among other studies, W2V has been employed to encode amino acid sequences, aiming to captivate hidden semantic properties of peptides through contextual word learning [26].

W2V provides two alternative model architectures, namely Continuous Bag of Words (CBOW) and Skip-Gram (SG). After conducting several experiments, we found that the SG model generally outperformed the CBOW. In terms of its underlying algorithm, W2V processes and stores successive sequences of k-mer amino acids within a specified window of the sequence, treating each sequence as a distinct unit and representing it with a numerical vector. Our W2V model was designed to encode a sequence window of n amino acids into an (n - k + 1) x S matrix, where S is the dimensionality of each vector representation.

We found that our model achieved better performance when we set the maximum amino acid sequence length (seqwin) to match the length of the longest peptide sequence among training datasets, i.e., the value 36, 61 and 27 for the CPP-Classification, Uptake-Efficiency and PMO-Delivery tasks, respectively. For the sequences with a length smaller than seqwin, we applied zero padding. Finally, regarding the implementation of W2V, we utilized the free open-source Python library Gensim [27]. We thoroughly examined the training parameters of W2V model, including the values of k-mer, vector size, number of training epochs, and window size (Additional File 1, Table S1).

Inspired by Kurata’s et al. [26] approach, we constructed three dataset-specific W2V dictionaries from the training and test datasets, one for each of the studied tasks, by adopting their sandwich structure concept. The training dataset is divided into subsets of positive and negative samples. Afterwards, the test dataset is shuffled and interleaved between them. This process resulted in our W2V dictionaries, from which our word embeddings were generated.

### Pre-trained LLMs approach

Approaching the task from a different perspective, we carefully selected three pre-trained models based on the LLMs: Text-To-Text Transfer Transformer (T5) [8], Bidirectional Encoder Representations from Transformers (BERT) [9], and Evolutionary Scale Modeling (ESM-2) [10], to generate the embeddings of amino acid sequences. Similar to the usage of non-protein-based LLMs in language-based tasks, these models have been pre-trained on large protein databases comprising millions of sequences, where each amino acid is treated as a word and each sequence as a sentence, demonstrating their ability to reveal underlying semantic characteristics of peptides. The selected models have been designed following the Masked Language Modeling pre-training approach, where a portion of the input tokens, 15% in our case, is randomly replaced with the special [MASK] token. Subsequently, the transformer architecture-based model is trained to predict the masked tokens, considering the surrounding unmasked tokens, and calculating the discrepancy between the predictions and the original targets. Once the loss has been computed, the model’s parameters are iteratively updated through backpropagation to minimize the loss function, resulting in meaningful embeddings [9]. Models’ hyperparameters are provided in Additional File 1, Table S2.

Regarding the T5 LLM, we utilized the ProtT5-XL-UniRef50 model [28], which has been pre-trained in a self-supervised manner on UniRef50, an extensive collection of 45 million protein sequences sourced from the UniProt database [29]. UniRef50 includes only sequences with less than 50% sequence similarity. Interestingly, T5 utilizes both encoder and decoder transformer models, whereas BERT exclusively adapts the encoder component.

In the case of the BERT LLM, we employed the ProtBERT model [28], which has been pre-trained similarly to ProtT5-XL-UniRef50 without any *a priori* human labelling. ProtBERT was mainly trained on UniRef100, a comprehensive dataset of 217 million protein sequences [29]. A significant distinction between ProtBERT and ProtT5-XL-UniRef50 models is that the former treats each involved sequence as a complete document without incorporating next sentence prediction [30].

Finally, regarding the ESM-2 LLM, we experimentally selected the *esm2 t36 3B UR50D* model [31], which has been pre-trained on UniRef50 and finetuned on various databases including UniProt [29] and Protein Data Bank [32]. Additionally, the Unsupervised Data Augmentation technique is often used to enrich their data. Similar to T5 and BERT models, ESM-2 incorporates the transformer architecture and typically employs only the encoder component. Notably, ESM-2 was specifically designed for protein sequence analysis, capturing the structural characteristics of peptides [33].

### Machine Learning methods

To solve our classification problem for identifying CPPs, we employed three widely used conventional ML methods: SVM, RF and GB. An SVM constructs a hyperplane that maximizes the margin of separation between positive and negative classes by utilizing kernel functions, aiming to minimize misclassification. On the other side, both RF and GB are ensemble techniques that build multiple decision trees. RF constructs the decision trees in parallel and combines them for the final prediction, while GB builds them sequentially, where each tree corrects the errors of its predecessor.

Apart from the three traditional ML algorithms, we employed three deep learning algorithms (DL), namely Convolutional Neural Network (CNN), Transformer (TX), and Bidirectional Long Short-Term Memory (bLSTM). We applied a CNN with two convolutional layers, to extract significant features, and two max-pooling layers, to reduce spatial dimensions while preserving meaningful features. From TX, we utilized the encoder component to transform the amino acid sequences, while we set the number of attention heads and the depth of the neural network to 3 and 4, respectively. In the case of bLSTM, in order to identify hidden patterns of peptides, we selected an improved version of a Recurrent Neural Network (RNN) with memory cells and a gating structure to avoid vanishing gradient problems, while we set the number of expected features in the input data equal to the vector size of the W2V model and the number of features in the hidden state to 128.

### Construction of ML-based models

Based on our encoding approach (W2V or Pre-trained LLM), we tested various models to identify the most promising for our studied tasks. We evaluated the performance of our models using both Cross-Validation (CV) and independent testing. Specifically, we implemented a 10-fold CV for the CPP-Classification, a Jackknife Validation for the Uptake-Efficiency and a 3-fold CV for the PMO-Delivery [2, 16].

For the W2V approach, except for testing W2V parameters, i.e., seqwin, vector size, epochs, sg and window size, we evaluated our models for k-mer values within the range of 1 to 10 [26]. Furthermore, for both of our encoding approaches, regarding RF, we executed the grid search that is proposed by Wolfe et al. [16] (Table 2), for the most significant parameters n estimators, max depth and max features. In the case of SVM, we tried to optimize the parameters C and Gamma, to handle the trade-off between classification error and margin, and to determine the impact of each training sample on the decision boundary, respectively.

**Table 2.**
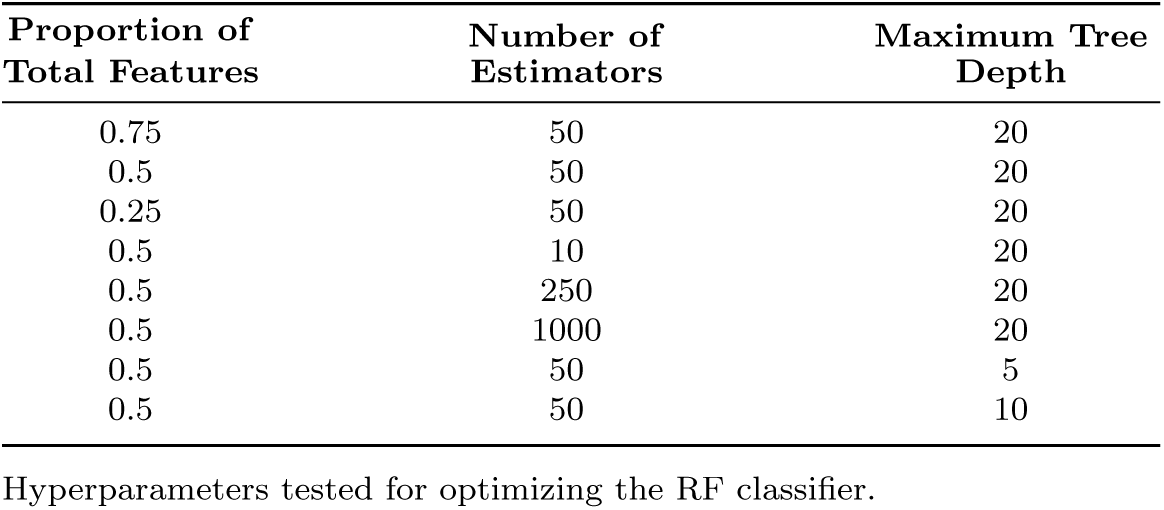
Grid search to optimize RF classifier’s hyperparameters.

More details about hyperparameter optimization for both ML and DL methods are provided in Additional File 1, Table S3. The SVM, RF and GB classifiers were implemented using scikit-learn, while the CNN, TX and bLSTM models were executed through PyTorch [34].

### Performance evaluation

To evaluate the prediction performance of the proposed models and compare them with state-of-the-art tools, we used six statistical measures, including Sensitivity (SE), Specificity (SP), Accuracy (ACC), Precision, Matthew’s Correlation Coefficient (MCC) and Area Under the Receiver Operating Characteristic Curve (AUC). The first five measures are calculated by the following equations:

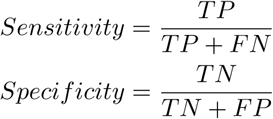

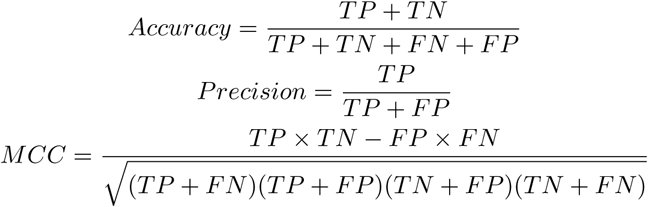

where TP, TN, FP, and FN denote the numbers of true positive, true negative, false positive and false negative, respectively. Finally, the AUC metric is computed by integrating the ROC curve, the graphical representation of SE against 1 – SP.

## Results

In this section, we outline the approach that we followed to select our proposed models and we present their overall performance across all tasks and datasets (Table 3). Furthermore, we include a detailed task-centered comparison of CPP2Vec against state-of-the art tools, including a visualized performance of both CPP2Vec and CPP2LLM on each task separately. Finally, we provide six radar charts to depict the overall predictive performance of each model across all datasets, highlighting their strengths and weaknesses (Figure 8).

**Table 3.** Performance metrics of CPP2Vec and CPP2LLM across tasks and datasets.

| Task | Dataset | SE | SP | ACC | MCC | AUC | Precision |
| --- | --- | --- | --- | --- | --- | --- | --- |
| **CPP2Vec** |  |  |  |  |  |  |  |
| **CPP Classification** | Validation | 0.845 | 0.840 | 0.842 | 0.687 | 0.915 | 0.842 |
|  | Test: *kelm* | 0.781 | 0.864 | 0.822 | 0.648 | 0.889 | 0.852 |
|  | Test: *mlcpp* | 0.906 | 0.646 | 0.776 | 0.572 | 0.877 | 0.719 |
| **Uptake Efficiency** | Validation | 0.754 | 0.679 | 0.716 | 0.434 | 0.727 | 0.701 |
|  | Test | 0.444 | 0.756 | 0.625 | 0.211 | 0.707 | 0.571 |
| **PMO Delivery** | Validation | 0.719 | 0.858 | 0.782 | 0.589 | 0.839 | 0.800 |
|  | Test | 0.800 | 1.000 | 0.857 | 0.730 | 1.000 | 1.000 |
| **CPP2LLM** |  |  |  |  |  |  |  |
| **CPP Classification** | Validation | 0.852 | 0.816 | 0.834 | 0.671 | 0.908 | 0.824 |
|  | Test: *kelm* | 0.714 | 0.877 | 0.795 | 0.599 | 0.832 | 0.853 |
|  | Test: *mlcpp* | 0.904 | 0.692 | 0.798 | 0.610 | 0.876 | 0.746 |
| **Uptake Efficiency** | Validation | 0.610 | 0.695 | 0.652 | 0.306 | 0.698 | 0.667 |
|  | Test | 0.552 | 0.731 | 0.656 | 0.288 | 0.662 | 0.600 |
| **PMO Delivery** | Validation | 0.700 | 0.889 | 0.781 | 0.624 | 0.890 | 0.861 |
|  | Test | 0.534 | 1.000 | 0.667 | 0.512 | 0.983 | 1.000 |

SE: Sensitivity, SP: Specificity, ACC: Accuracy, MCC: Matthews Correlation Coefficient, AUC: Area Under Curve.

### Selection of CPP2Vec and CPP2LLM

#### CPP2Vec

To determine which W2V and ML model provides the best results for each task, we executed a series of experiments, aiming to find the optimal value for each one of the involved parameters. Selected results are provided in Additional File 2. In the first step, we focused on the W2V model optimization. We started by stabilizing the ML model and experimenting with vector size, epochs, sg and window parameters. For instance, in Table 4 we provide a selection of representative results from our experiments on W2V model hyperparameter optimization for the PMO-Delivery task. We worked in a similar way for each one of the three studied tasks.

**Table 4.** Selected representative results from our W2V model hyperparameter optimization experiments for the PMO-Delivery task.

| W2V Parameters | Validation |  | Test |  |
| --- | --- | --- | --- | --- |
|  | ACC | MCC | ACC | MCC |
| 27, 300, 8, 1, 20 | 0.748 | 0.502 | 0.619 | 0.449 |
| 27, 100, 12, 1, 20 | 0.780 | 0.549 | 0.619 | 0.449 |
| 27, 100, 20, 0, 20 | 0.670 | 0.323 | 0.571 | 0.387 |
| 27, 100, 20, 1, 10 | 0.766 | 0.523 | 0.571 | 0.401 |
| **27, 100, 20, 1, 20** | **0.782** | **0.560** | **0.857** | **0.676** |

W2V Parameters are depicted as: (seqwin, size, epochs, sg, window). ACC: Accuracy, MCC: Matthews Correlation Coefficient.

Subsequently, we attempted to identify the most promising ML algorithms. For the CPP-Classification task we tested various ML algorithms, including RF, SVC, GB, TX, bLSTM and CNN. In Table 5 we present some selected results, that led us to select SVC with C=1.0 as the optimal model.

**Table 5.** Performance metrics of ML Models across validation and test datasets.

| ML Model<br>Parameters | Validation |  | Test: *kelm* |  | Test: *mlcpp* |  |
| --- | --- | --- | --- | --- | --- | --- |
|  | ACC | MCC | ACC | MCC | ACC | MCC |
| RF | 0.783 | 0.567 | 0.775 | 0.551 | 0.729 | 0.476 |
| GB | 0.811 | 0.624 | 0.791 | 0.582 | 0.768 | 0.538 |
| TX | 0.834 | 0.671 | **0.826** | **0.654** | 0.756 | 0.516 |
| LSTM | 0.808 | 0.621 | 0.782 | 0.567 | 0.725 | 0.462 |
| CNN | 0.819 | 0.644 | 0.797 | 0.597 | 0.739 | 0.487 |
| **SVC** | **0.842** | **0.687** | 0.822 | 0.648 | **0.776** | **0.572** |

RF(max\_depth=4, n\_estimators=10), SVC(C=1.0, probability=True), GB(n\_estimators=100, learning\_rate=0.1, max\_depth=4).

For the Uptake-Efficiency task, we executed some initial trials with SVC models, based on the CPP-Classification results. We conducted the grid search proposed by Wolfe et al. [16] (Table 2) and we compared the results with the SVCs. We observed that SVC with C=10.0 outperforms best RF model with validation accuracy 71.7% compared to 67.9%, respectively, leading us to its selection.

Finally, as the accuracy of the Uptake-Efficiency task model remained low even after optimizing the model’s parameters, we decided to use the 64-peptides dataset as a training dataset to predict PMO-Delivery efficiency. We initially found the best values for the W2V model and then we executed the same grid search. The proposed W2V model values for CPP-Classification, Uptake-Efficiency and PMO-Delivery tasks are depicted in Table 6.

**Table 6.** Overview of the proposed W2V model parameters per task.

| W2V<br>Parameter | CPP<br>Classification | Uptake<br>Efficiency | PMO<br>Delivery |
| --- | --- | --- | --- |
| kmer | 2 | 3 | 1 |
| seqwin | 36 | 61 | 27 |
| vector size | 200 | 300 | 100 |
| epochs | 8 | 12 | 20 |
| sg | 1 | 1 | 1 |
| window | 20 | 20 | 20 |

Values correspond to the optimal W2V hyperparameters selected per task.

The only difference is that in this case we repeated our experiments for the best two models, as datasets consist of a small number of sequences. The average metrics for both StratifiedKFold and KFold cross-validators [35] can be found in the Additional File 1, Tables S4 and S5. The proposed ML models for all tasks are depicted at Table 7.

**Table 7.** Proposed ML models per task of CPP2Vec.

| Task | ML Model |
| --- | --- |
| CPP-Classification | SVC (C=1.0, prob=True) |
| Uptake-Efficiency | SVC (C=10.0, prob=True) |
| PMO-Delivery | RF (max_features=0.5, max_depth=20, n_estimators=250) |

RF in (max\_features, max\_depth, n\_estimators) format.

#### CPP2LLM

For each of the tasks we studied, i.e. CPP-Classification, Uptake-Efficiency, and PMO-Delivery, we conducted 12 experiments per LLM (T5, BERT and ESM-2). Specifically, for each task and each LLM we ran 3 experiments with SVC (C=1.0, C=5.0 and C=10.0), 1 experiment with GB (n estimators=100, learning rate=0.1, max depth=4) and 8 experiments with the RF classifiers of the proposed grid search. By taking into consideration the results and their running times we selected the LLM and ML models that are presented in Table 8.

**Table 8.** Proposed LLM and ML models per task of CPP2LLM.

| Task | LLM Model | ML Model |
| --- | --- | --- |
| CPP-Classification | BERT | RF (max_features=0.5, max_depth=20, n_estimators=1000) |
| Uptake-Efficiency | ESM-2 | SVC (C=10.0, prob=True) |
| PMO-Delivery | ESM-2 | RF (max_features=0.25, max_depth=20, n_estimators=50) |

RF in (max\_features, max\_depth, n\_estimators) format.

#### Visualization of CPP2Vec embeddings

To validate the effectiveness of our proposed models in distinguishing between the studied classes, i.e. CPP/non-CPP, High/Low uptake efficiency, and *≥*3-fold/*<*3-fold improvement, we used Uniform Manifold Approximation and Projection (UMAP) [36] and Principal Component Analysis (PCA) [37] to visualize the learned W2V embeddings in 2D space.

In Figure 2 we present the UMAP plot for the *kelm* dataset, where distinct clusters for each class are depicted, indicating the model’s ability to successfully capture meaningful patterns in the data. The respective plots for the *mlcpp*, 64-peptides, and 7-novel-sequences datasets can be found in Additional File 1, specifically in Figures S1, S2, and S3, respectively, where well-defined boundaries between clusters validate the classification performance of CPP2Vec.

**Fig. 2.**
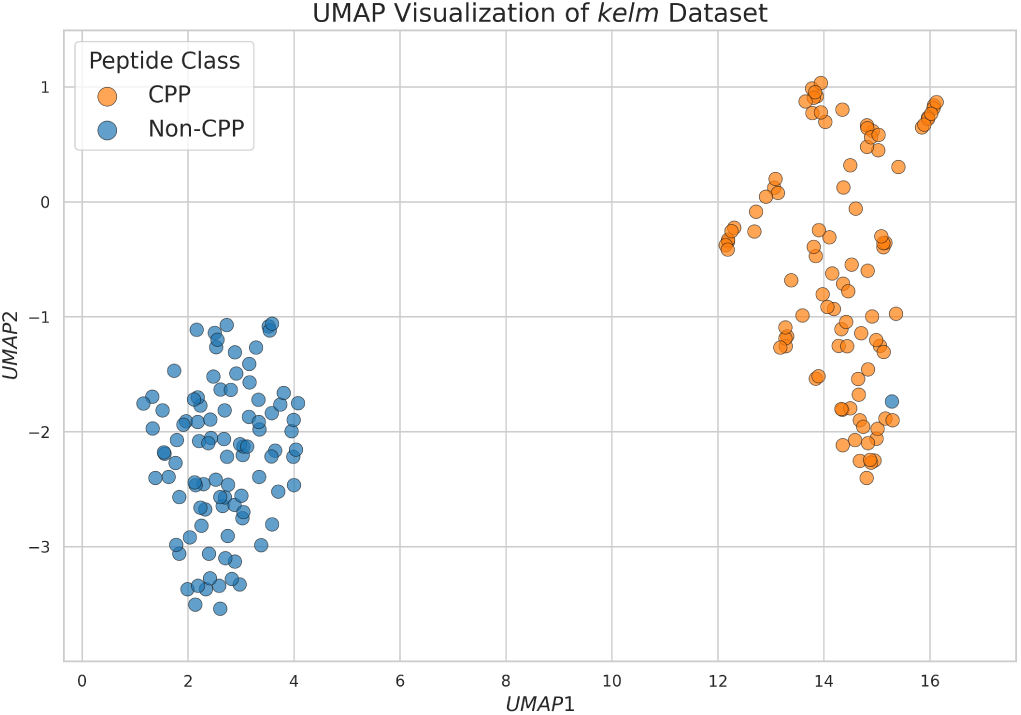
UMAP visualization of the *kelm* dataset. CPPs and non-CPPs are depicted in orange and blue, respectively. The clear distinction between the two classes highlights the representation power of CPP2Vec.

In Figure 3 we present the heatmap of the first five PCA components across peptides, grouped by class labels and sorted in descending predicted probability scores, for the *mlcpp* dataset. Differences in color intensity between classes suggest that these PCA components can effectively distinguish the peptides. Within each class, the existence of color patterns among certain peptides reveals the possibility that these peptides share common characteristics, while some PCA components show greater variability, indicating that there are features that contribute less heavily to class distinction. The heatmaps for the other test datasets are depicted in Additional File 1, Figures S4, S5, and S6.

**Fig. 3.**
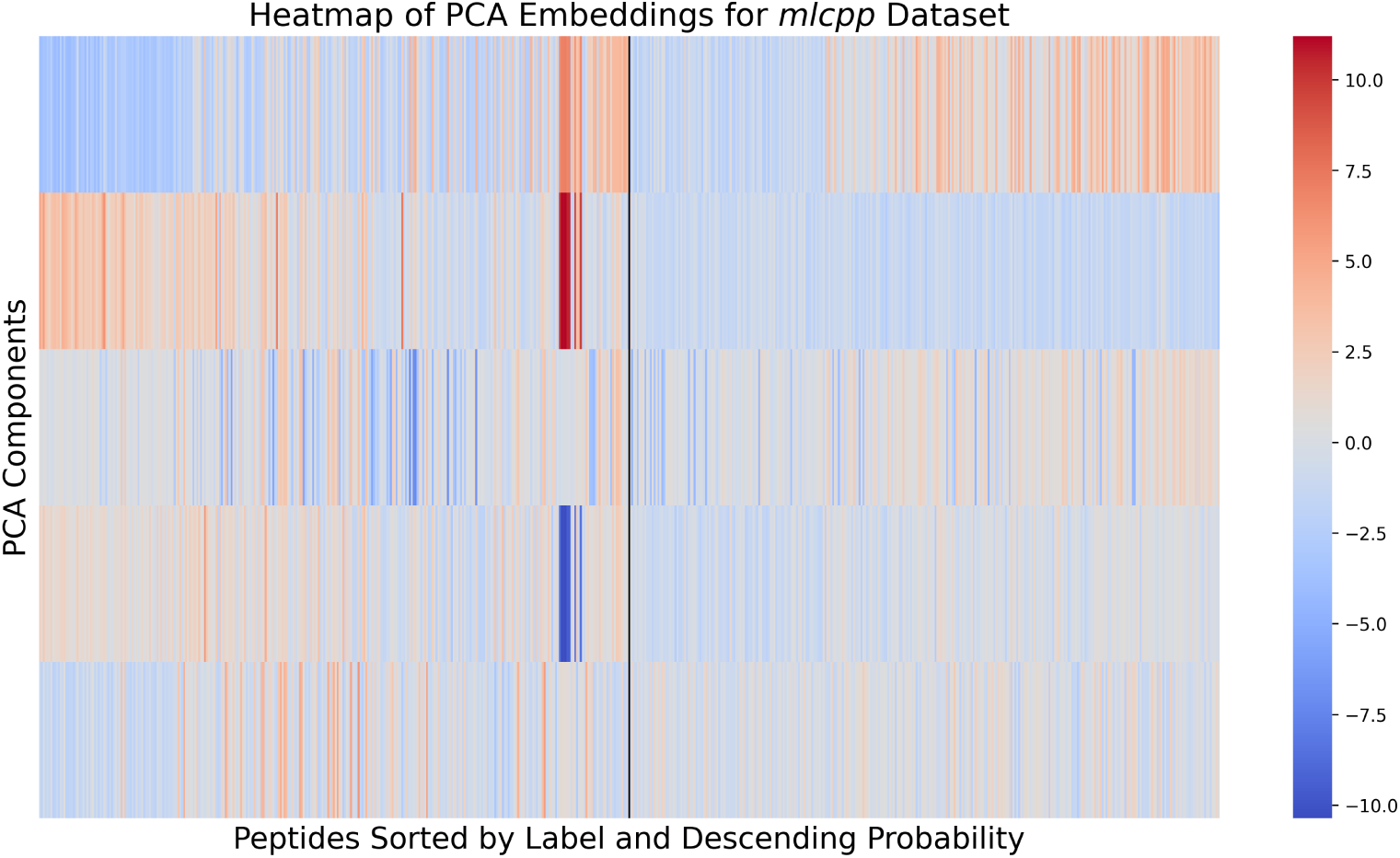
Heatmap of the first five PCA Components for the *mlcpp* dataset. Peptides are grouped by labels from CPPs to Non-CPPs and sorted in descending predicted probability scores. The black vertical line illustrates the boundary between classes.

#### Comparison of CPP2Vec with state-of-the-art tools

##### CPP-Classification

To evaluate the performance of CPP2Vec against state-of-the-art tools at the CPP-Classification task, we conducted experiments using the same datasets employed in the comparative study by Su et al. [2] A summary of the results among selected models can be seen in Figures 4 and 5, for the *kelm* and *mlcpp* datasets, respectively, while a full comparison with all the available models can be found in Additional File 1, Tables S6, S7.

**Fig. 4.**
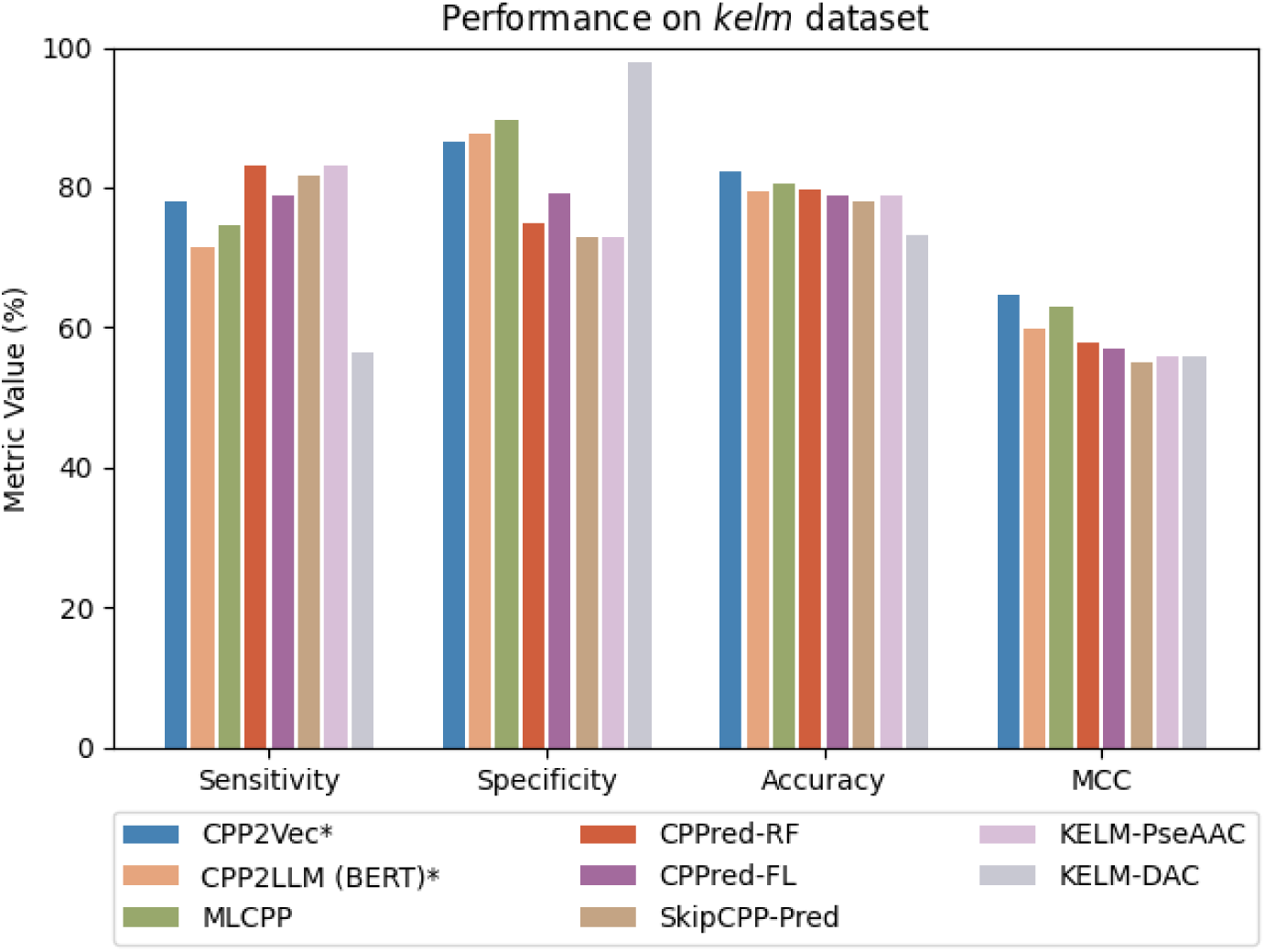
Comparison of CPP2Vec and CPP2LLM with state-of-the-art tools on *kelm* dataset. Our proposed models are marked with an asterisk (*). Each bar represents the mean performance across the 10-fold CV. The selected protein-based LLM of CPP2LLM is shown in parentheses.

**Fig. 5.**
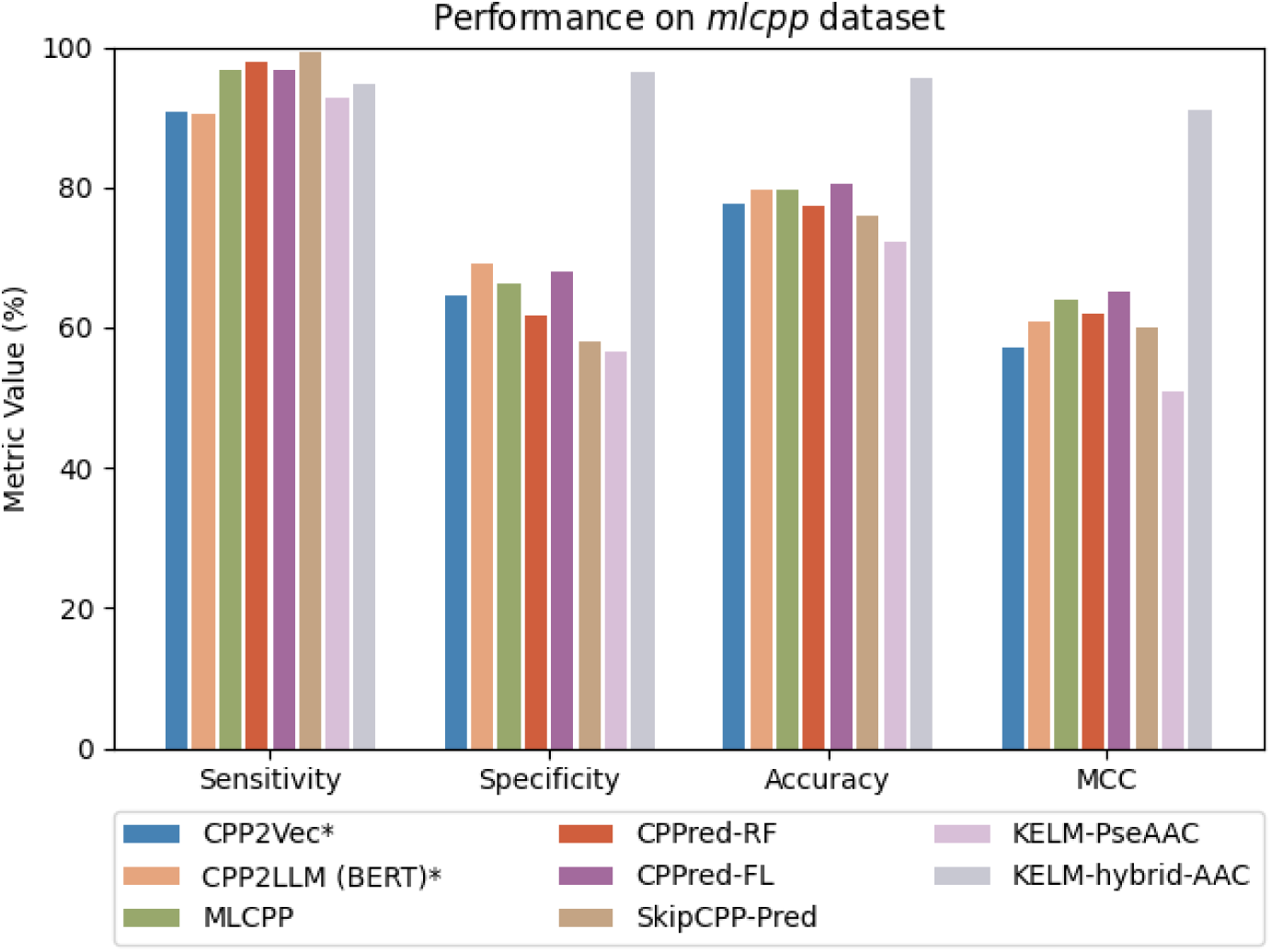
Comparison of CPP2Vec and CPP2LLM with state-of-the-art tools on *mlcpp* dataset. Our proposed models are marked with an asterisk (*). Each bar represents the mean performance across the 10-fold CV. The selected protein-based LLM of CPP2LLM is shown in parentheses.

On the *kelm*, CPP2Vec consistently outperforms many state-of-the-art tools. In particular, CPP2Vec achieved the highest overall ACC at 82.29%, compared to the closest tool, MLCPP, which reached 80.67%, while CPPred-RF and KELM-AAC reported lower accuracies of 79.83% and 77.31%, respectively. CPP2Vec also achieved the highest MCC score of 0.65, outperforming MLCPP (0.63) and other tools like CPPred-RF (0.58) and SkipCPP-Pred (0.55), highlighting its superior balance between true positives and false positives. Additionally, CPP2Vec reached a SP of 86.45% with a SE of 78.12%. Although some models demonstrated higher SP, such as KELM-DAC (97.92%) and CellPPD (93.75%), their trade-off in SE is noteworthy, with both models only achieving 56.34% and 63.38%, respectively.

On the *mlcpp*, while CPP2Vec does not reach the highest metrics (ACC: 95.61%, MCC: 0.91, SE: 94.63%, SP: 96.37%) of the KELM-hybrid-AAC, it still demonstrates competitive performance against state-of-the-art tools. CPP2Vec achieves an ACC of 77.65%, comparable to models MLCPP (79.53%), CPPred-RF (77.49%), CellPPD (77.78%), and SkipCPP-Pred (76.02%), while outperforming some of the KELM-CPPpred methods, including KELM-hybrid-PseAAC (72.22%) and KELM-DAC (71.35%). In terms of MCC, CPP2Vec achieves a moderate score of 0.57, outperforming KELM-hybrid-PseAAC (0.51) and KELM-DAC (0.50), while reamaining comparable to CellPPD (0.56) and SkipCPP-Pred (0.60), though falling short of MLCPP (0.64) and CPPred-RF (0.62). Interestingly, CPP2vec a high SE of 90.67%, while also maintains a SP of 64.63%, which is higher than KELM-hybrid-PseAAC (56.48%), KELM-DAC (54.92%), SkipCPP-Pred (58.03%), and CPPred-RF (61.66%), and comparable to MLCPP (66.32%) and CPPred-FL (67.88%). This more balanced SE and SP indicates that CPP2Vec can credibly detect CPPs without sacrificing its ability to filter out false positives.

Consequently, CPP2Vec demonstrates consistently high performance in the CPP-Classification task, maintaining strong and reliable results across both the *kelm* and *mlcpp* datasets, without requiring any feature engineering. This avoids the risk of overperformance due to biased, dataset-specific conditions.

##### Uptake-Efficiency

To evaluate the performance of CPP2Vec at the Uptake-Efficiency task, we compared the results of jackknife validation with the metrics of state-of-the-art predictors derived from Manavalan’s et al. [15] study, on the CPPsite3 dataset [7] (Figure 6).

**Fig. 6.**
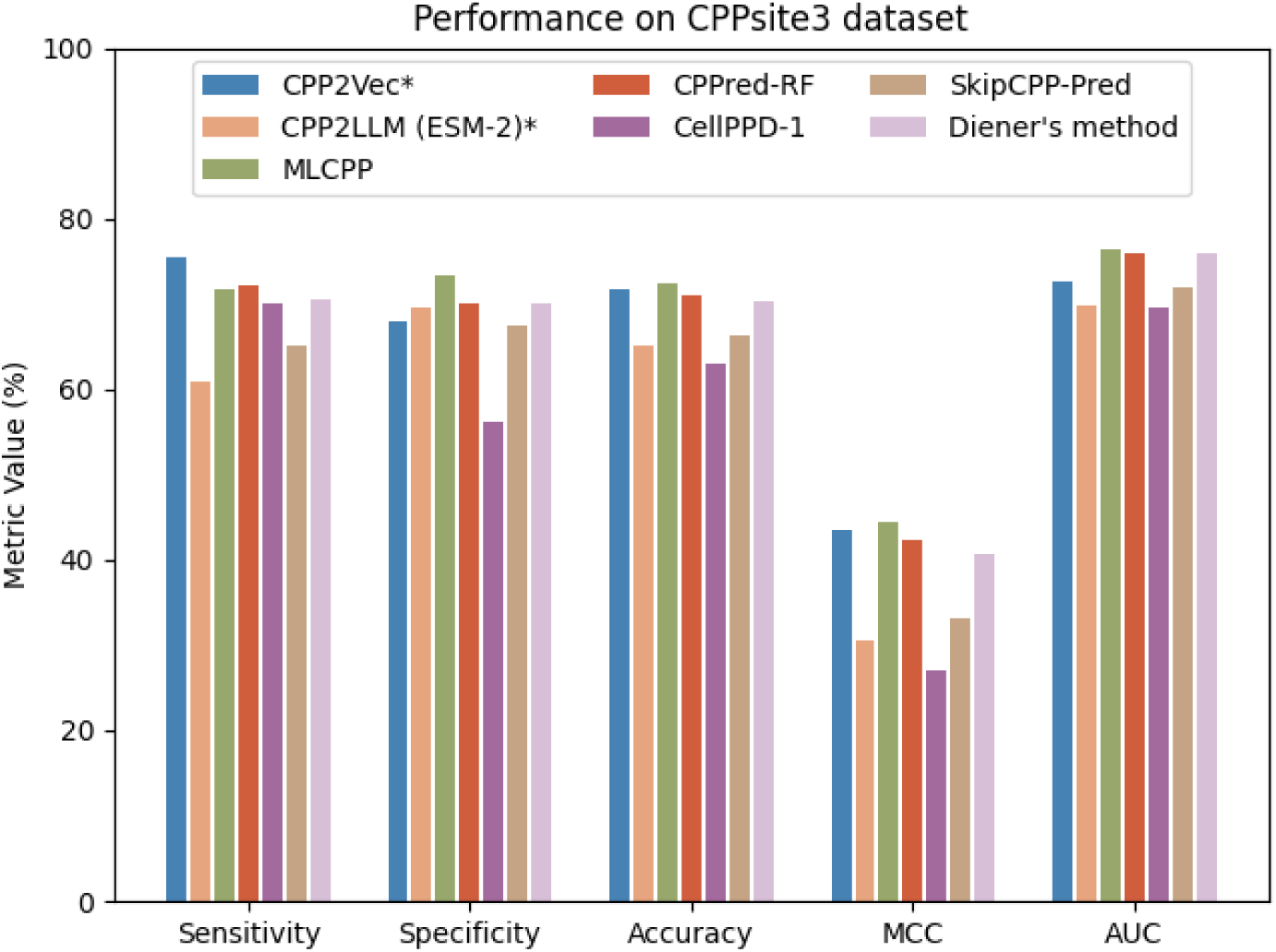
Comparison of CPP2Vec and CPP2LLM with state-of-the-art tools on CPPsite3 dataset. Our proposed models are marked with an asterisk (*). Each bar represents the mean performance across the Jackknife Validation. The selected protein-based LLM of CPP2LLM is shown in parentheses.

CPP2Vec scores the highest SE of 75.4% among all the compared methods, highlighting its ability to accurately detect peptides with high uptake efficiency. In terms of overall ACC, CPP2Vec scores 71.6%, closely trailing MLCPP (72.5%), while out-performing other models such as SkipCPP-Pred (66.3%), CellPPD (63.1%), and CPPred-RF (71.1%). For MCC, CPP2Vec achieves the second-best performance with a score of 0.434, following MLCPP (0.445), and ahead of CPPred-RF (0.423), Diener’s method (0.406), SkipCPP-Pred (0.332), and CellPPD (0.270). Finally, CPP2Vec reaches an AUC of 72.7% and a SP of 67.9%, keeping it in a competitive range, although MLCPP exceeds it with an AUC of 76.4% and a SP of 73.3%.

##### PMO-Delivery

As none of the current state-of-the-art CPP prediction tools provide a model for detecting peptides that could enhance PMO-Delivery across the plasma membrane, we compared the performance of CPP2Vec to the method that Wolfe et al. [16] proposed in their study, on the 64-peptides dataset (Figure 7). Specifically, CPP2Vec showed an accuracy of 78%, a precision of 80% and a recall of 72%, which are 6%, 5% and 3% higher than Wolfe’s et al. method, respectively.

**Fig. 7.**
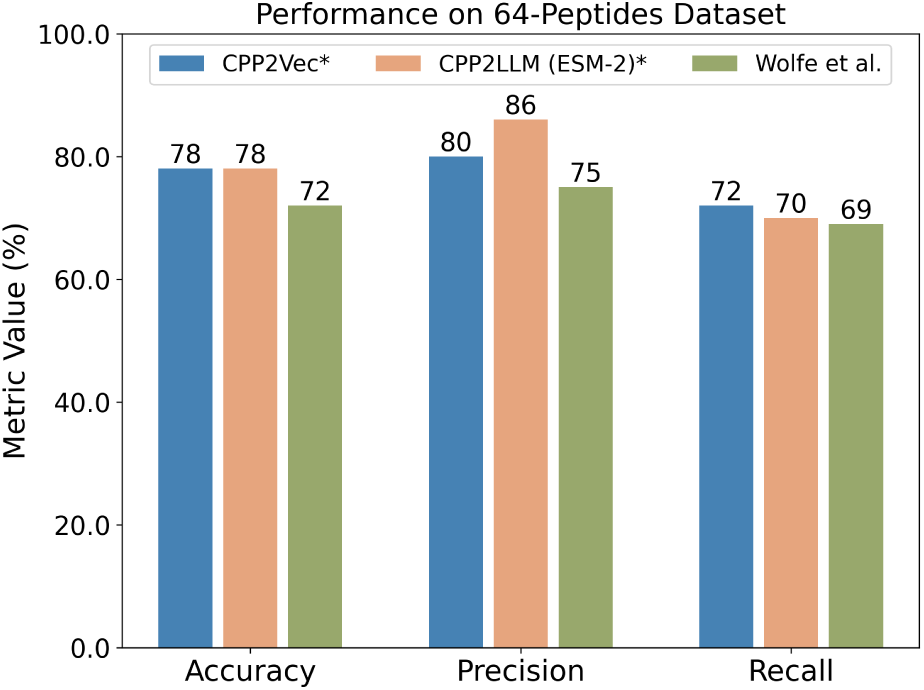
Comparison of CPP2Vec and CPP2LLM with Wolfe’s et al. [16] method on 64-peptides dataset. Our proposed models are marked with an asterisk (*). Each bar represents the mean performance across the 3-fold CV. The selected protein-based LLM of CPP2LLM is shown in parentheses.

**Fig. 8.**
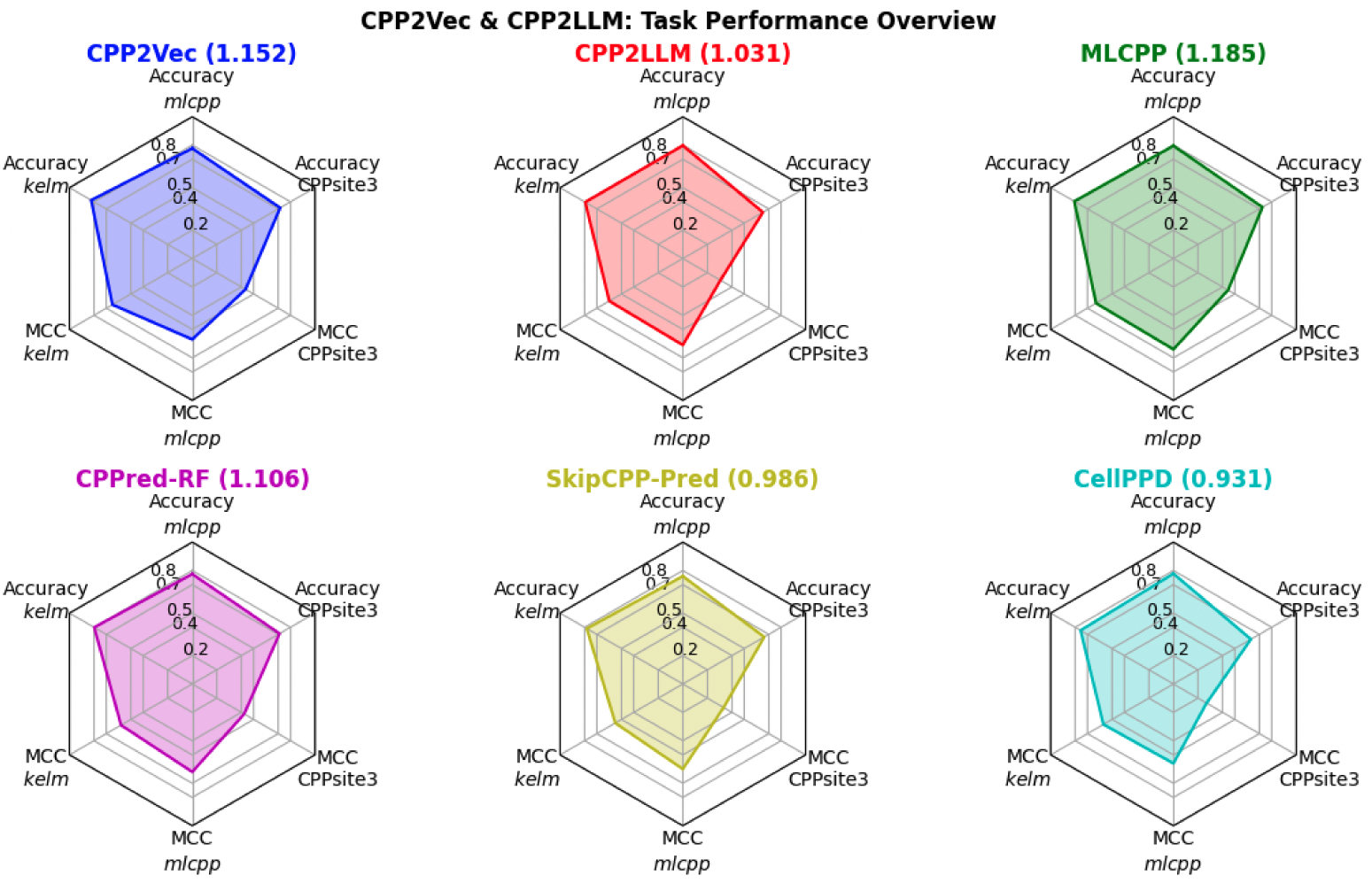
Depiction of the overall predictive performance per model. Each polygon represents a tool, i.e. MLCPP, CPPred-RF, SkipCPP-Pred, CellPPD, as well as our proposed models, CPP2Vec and CPP2LLM. The top three edges represent the accuracy of the models on the *kelm*, *mlcpp*, and CPPsite3 datasets, while the bottom edges denote their respective MCC scores. In general, larger areas indicate better performance (values in parentheses).

##### Overall Predictive Performance

Figure 8 illustrates the predictive performance of each model across the CPP-Classification and Uptake-Efficiency tasks. We present six radar charts where each polygon corresponds to a specific tool, including MLCPP, CPPred-RF, SkipCPP-Pred, CellPPD, as well as our proposed models, CPP2Vec and CPP2LLM. The top three edges of each polygon represent the accuracy of the models on the *kelm*, *mlcpp*, and CPPsite3 datasets, while the bottom three edges denote their respective MCC scores.

Even though CPP2Vec shows slightly weaker results in a few instances compared to the state-of-the-art tools, it consistently provides more stable outcomes overall. Notably, CPP2Vec stands out for its balanced and relatively high performance across all datasets, demonstrating a strong ability to generalize across diverse predictive scenarios. This consistency across different tasks highlights its reliability in the field of CPP prediction.

##### Case Study

We selected manually from the available literature 14 peptides that have been tested for their PMO-Delivery efficiency on mdx mice [18]. The mdx mouse is a famous model that was discovered in 1984 and it has been extensively used to study DMD [12, 13]. The mdx mouse presents a point mutation at exon 23 of its DMD gene, where a Guanine changes to a Thymine causing a premature stop codon, which leads to the production of a small non-functional dystrophin protein. The symptoms are not as severe as in humans; however, they share a lot of similarities in their DMD genes making them a proper model for *in vitro* and *in vivo* trials.

Details about the 14 selected peptides are provided in Table 9. The first four peptides, namely muscle-specific peptide (MSP), trans-activating transcriptional activator (TAT), adeno-associated virus 6 (AAV6) and AAV8, have been used by Yin et al. [38] in 2007, administering a single intramuscular injection (5*µ*g IM) in the tibialis anterior (TA) muscles of 2-month-old mdx mice. Interestingly in their study they used PNAs instead of PMOs, for stronger stability. In 2008, Matthew Wood et al. [39] tested naked B-peptide and connected it to PMO, with a single 5*µ*g IM at TA muscle with promising results, too. Finally, in 2008 Jearawiriyapaisarn et al. [40] constructed a series of peptides with improved, not only muscle, but also cardiac exon skipping activity for DMD treatment [41]. This was the first study where exon skipping was detected in cardiac muscle, reaching 15% of normal levels of dystrophin protein.

**Table 9.**
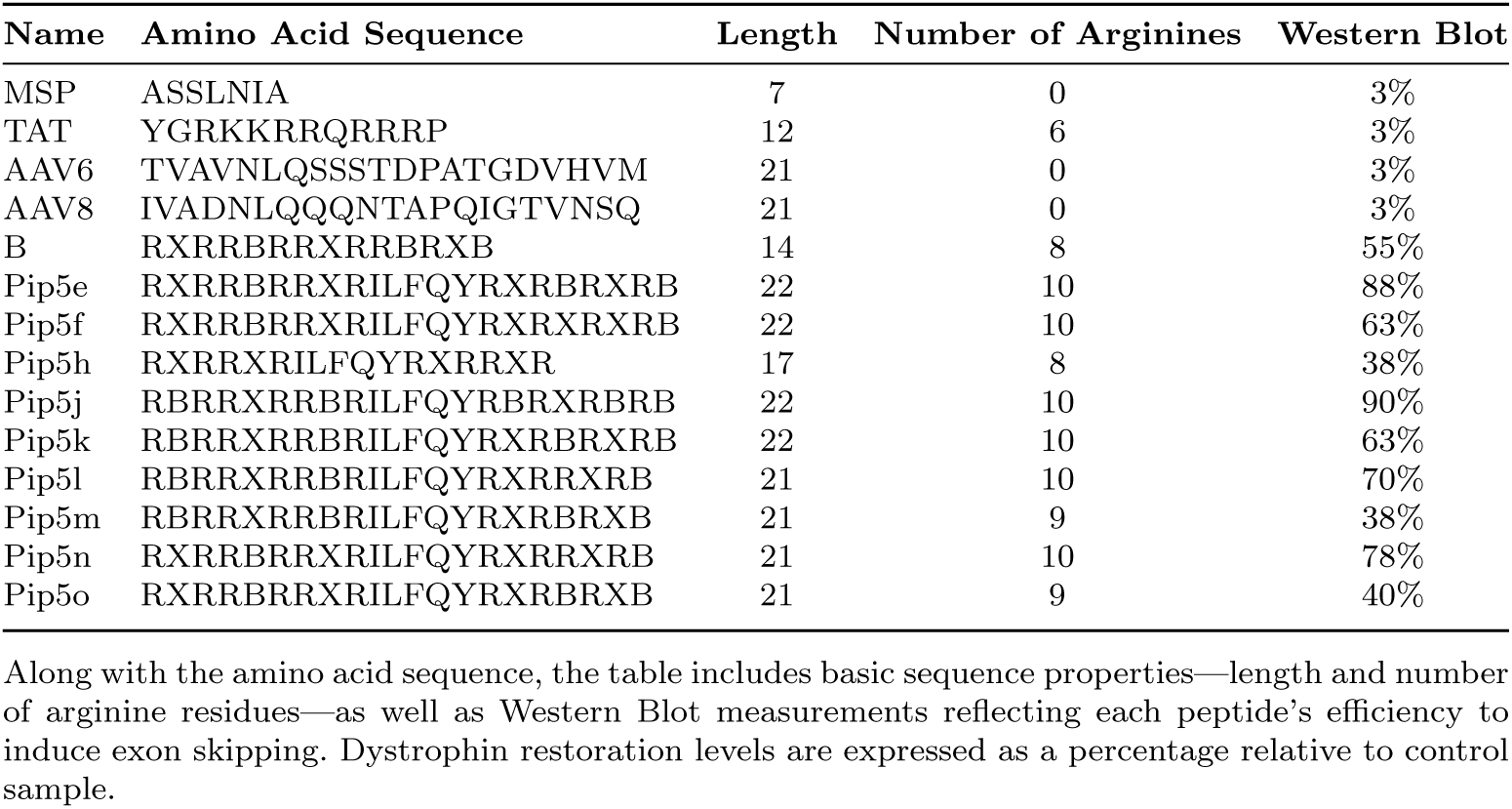
Amino acid sequences and properties of the 14 peptides of the DMD dataset.

We substituted *β*-alanine (B) with *α*-alanine (A) and 6-aminohexanoic acid (X) with lysine (K), respectively, to avoid non-natural amino acids for fair comparisons.

Finally, we tested our PMO-Delivery model on the DMD dataset, to thoroughly evaluate its performance, assessing its practical applicability and effectiveness in handling real case challenges. We calculated the SCC and PCC between the predicted probabilities and the Western Blot percentages, and we depicted the model’s performance using a Scatter Plot (Figure 9). We clearly delineated between efficient and inefficient peptides, highlighting the ability of CPP2Vec to accurately predict peptide delivery efficacy.

**Fig. 9.**
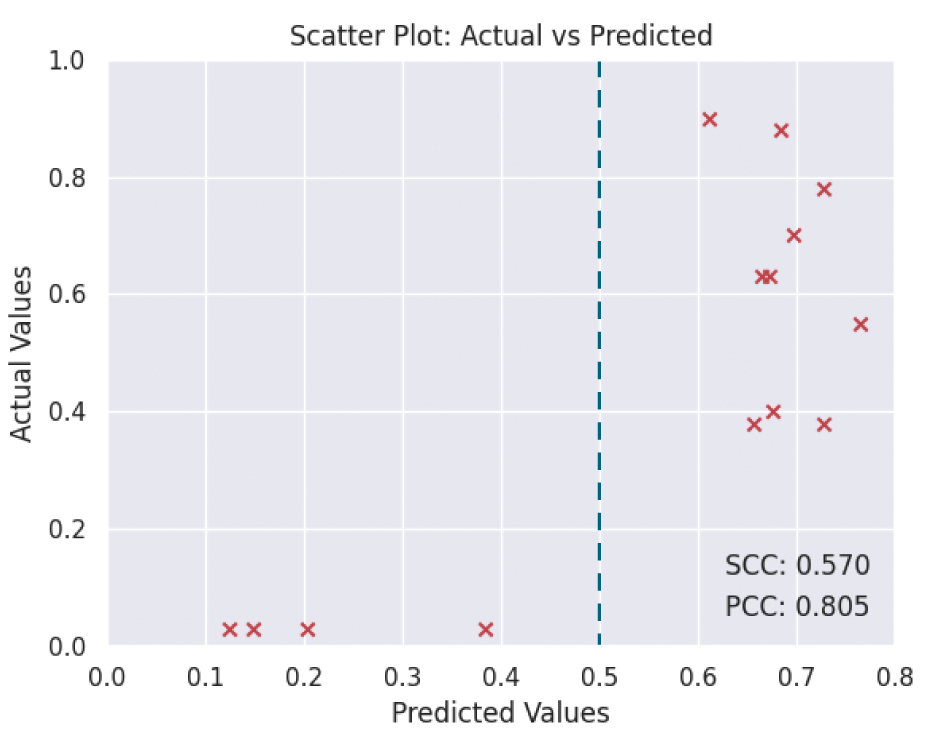
Scatter plot of the predicted vs actual values on the DMD dataset. Each red cross represents one of the 14 peptides. The vertical dashed line on 0.5 illustrates the model’s ability to accurately distinguish efficient peptides from inefficient ones.

## Discussion

In this study, we introduced CPP2Vec, an ML-based CPP prediction tool that utilizes the W2V technique to generate the amino acid sequence embeddings, aiming to capture any hidden patterns of peptides. Given the high cost of wet-lab experiments, in silico methods like CPP2Vec, provide an efficient approach to detect novel CPPs and categorize them based on their uptake efficiency. To achieve this, we developed three ML classifiers, including experiments with DL models.

Our proposed models were extensively evaluated and compared against state-of-the-art tools, achieving significant predictive performance. A key advantage of CPP2Vec is its ability to deliver accurate and robust predictions without requiring any prior feature engineering, relying only on the amino acid sequence of peptides. As expected, we confirmed that RF and SVC models outperformed DL models, likely due to the limited size of relevant datasets. Furthermore, similar to other state-of-the-art tools, CPP2Vec’s performance on the Uptake-Efficiency task remains moderate, which can be associated with the unclear internalization mechanisms of CPPs and the insufficient amount of data.

Approaching the CPP prediction task from a different angle, we explored using protein-based LLMs to generate contextualized embeddings - an approach not previously applied in the CPP prediction field, similar to W2V. Although this methodology produced results comparable to CPP2Vec, it required significantly more computational resources, which made CPP2Vec a better choice overall in terms of performance, computational requirements, and runtime.

In addition, we introduced a specialized PMO-Delivery model designed to predict whether a peptide could enhance the delivery of a PMO-complex into the cell compared to its naked version. To investigate the generalization ability of our model across tasks, we executed a case study on DMD, where its robust performance highlighted its potential for discovering novel CPPs.

## Conclusion

In this research, we developed CPP2Vec, an ML-based tool for CPP prediction that utilizes the W2V approach to encode amino acid sequences, aiming to uncover underlying peptide patterns. We propose three ML models, each tailored to excel in one of the following tasks: CPP Classification, Uptake Efficiency and PMO Delivery. A key advantage of CPP2Vec is that is designed to eliminate the need for manual, task-specific feature engineering, by automatically learning meaningful representations from the data.

A comparison with state-of-the-art models, along with a case study focused on DMD, highlights CPP2Vec’s strong predictive performance. Based on its architecture and consistent results, we anticipate that CPP2Vec can serve as a valuable tool for experts across diverse therapeutic scenarios - especially considering the significant limitations posed by the high cost of wet-lab experiments.

## Abbreviations

*CPPs*: Cell-Penetrating Peptides
*ASOs*: Antisense Oligonucleotides
*PNAs*: Peptide Nucleic Acids
*PMOs*: Phosphorodiamidate Morpholino Oligomers
*DMD*: Duchenne Muscular Dystrophy
*LLMs*: Large Language Models
*ML*: Machine Learning
*AAC*: Amino Acid Composition
*CKS*: Composition of K-spaced Amino Acid Pairs
*PseAAC*: Pseudo Amino Acid Composition
*W2V*: Word2Vec
*SVM*: Support Vector Machine
*RF*: Random Forest
*GB*: Gradient Boosting
*NLP*: Natural Language Processing
*CBOW*: Continuous Bag of Words
*SG*: Skip-Gram
*T5*: Text-To-Text Transfer Transformer
*BERT*: Bidirectional Encoder Representations from Transformers
*ESM-2*: Evolutionary Scale Modeling - 2
*DL*: Deep Learning
*CNN*: Convolutional Neural Network
*TX*: Transformer
*bLSTM*: Bidirectional Long Short-Term Memory
*RNN*: Recurrent Neural Network
*CV*: Cross Validation
*SE*: Sensitivity
*SP*: Specificity
*ACC*: Accuracy
*MCC*: Matthew’s Correlation Coefficient
*AUC*: Area Under the Receiver Operating Characteristic Curve
*TP*: True Positive
*TN*: True Negative
*FP*: False Positive
*FN*: False Negative
*UMAP*: Uniform Manifold Approximation and Projection
*PCA*: Principal Component Analysis
*MSP*: Muscle-Specific Peptide
*TAT*: Trans-Activating Transcriptional Activator
*AAV*: Adeno-Associated Virus
*TA*: Tibialis Anterior

## Supporting Information

Additional File 1.

Additional File 2.

## Declarations

## Availability of data and material

CPP2Vec’s models along with the training and test datasets are available for use at: https://github.com/SSvolou/CPP2Vec.

## Authors’ Contributions

G.P., A.K., and V.K. conceived the initial idea of this study.; V.K. and A.K. supervised the research execution.; S.S. and V.K. designed the methodology and planned the experiments.; S.S. analyzed the data, implemented the computer code and conducted the experiments with contributions from V.K.; S.S. and V.K. collaborated on the interpretation of the results, incorporating feedback from A.K. and G.P.; S.S. wrote the original manuscript, receiving critical comments from V.K.; All authors reviewed the manuscript.

## References

[1] Xie J, Bi Y, Zhang H, Dong S, Teng L, Lee RJ, et al. Cell-penetrating peptides in diagnosis and treatment of human diseases: from preclinical research to clinical application. Frontiers in pharmacology. 2020;11:697.

[2] Su R, Hu J, Zou Q, Manavalan B, Wei L. Empirical comparison and analysis of web-based cell-penetrating peptide prediction tools. Briefings in bioinformatics. 2020;21(2):408–420.

[3] Pandey P, Patel V, George NV, Mallajosyula SS. KELM-CPPpred: Kernel extreme learning machine based prediction model for cell-penetrating peptides. Journal of proteome research. 2018;17(9):3214–3222.

[4] Wei L, Tang J, Zou Q. SkipCPP-Pred: an improved and promising sequence-based predictor for predicting cell-penetrating peptides. BMC genomics. 2017;18:1–11.

[5] Wei L, Xing P, Su R, Shi G, Ma ZS, Zou Q. CPPred-RF: a sequence-based predictor for identifying cell-penetrating peptides and their uptake efficiency. Journal of Proteome Research. 2017;16(5):2044–2053.

[6] Qiang X, Zhou C, Ye X, Du Pf, Su R, Wei L. CPPred-FL: a sequence-based predictor for large-scale identification of cell-penetrating peptides by feature representation learning. Briefings in Bioinformatics. 2020;21(1):11–23.

[7] Gautam A, Chaudhary K, Kumar R, Sharma A, Kapoor P, Tyagi A, et al. In silico approaches for designing highly effective cell penetrating peptides. Journal of translational medicine. 2013;11:1–12.

[8] Raffel C, Shazeer N, Roberts A, Lee K, Narang S, Matena M, et al. Exploring the limits of transfer learning with a unified text-to-text transformer. Journal of machine learning research. 2020;21(140):1–67.

[9] Devlin J. Bert: Pre-training of deep bidirectional transformers for language understanding. arXiv preprint arXiv:181004805. 2018;.

[10] Lin Z, Akin H, Rao R, Hie B, Zhu Z, Lu W, et al. Evolutionary-scale prediction of atomic-level protein structure with a language model. Science. 2023;379(6637):1123–1130.

[11] Mikolov T, Chen K, Corrado G, Dean J. Efficient estimation of word representations in vector space. arXiv preprint arXiv:13013781. 2013;3781.

[12] of Neurological Diseases NI, Stroke. Muscular Dystrophy: Hope Through Research. 72-77. Information Office, National Institute of Neurological Diseases and Stroke; 1971.

[13] Schultz TI, Raucci Jr FJ, Salloum FN. Cardiovascular disease in Duchenne muscular dystrophy: overview and insight into novel therapeutic targets. Basic to Translational Science. 2022;7(6):608–625.

[14] Diener C, Garza Ramos Martónez G, Moreno Blas D, Castillo Gonzalez DA, Corzo G, Castro-Obregon S, et al. Effective design of multifunctional peptides by combining compatible functions. PLoS Computational Biology. 2016;12(4):e1004786.

[15] Manavalan B, Subramaniyam S, Shin TH, Kim MO, Lee G. Machine-learning-based prediction of cell-penetrating peptides and their uptake efficiency with improved accuracy. Journal of proteome research. 2018;17(8):2715–2726.

[16] Wolfe JM, Fadzen CM, Choo ZN, Holden RL, Yao M, Hanson GJ, et al. Machine learning to predict cell-penetrating peptides for antisense delivery. ACS central science. 2018;4(4):512–520.

[17] Manavalan B, Patra MC. MLCPP 2.0: an updated cell-penetrating pep-tides and their uptake efficiency predictor. Journal of Molecular Biology. 2022;434(11):167604.

[18] Triantafyllidou V. Computational methods for optimization of therapeutic approaches for Duchenne Muscular Dystrophy [Master’s thesis]. National and Kapodistrian University of kAthens. Greece; 2018.

[19] Kardani K, Bolhassani A. Cppsite 2.0: An available database of experimentally validated cell-penetrating peptides predicting their secondary and tertiary structures. Journal of molecular biology. 2021;433(11):166703.

[20] Waghu FH, Barai RS, Gurung P, Idicula-Thomas S. CAMPR3: a database on sequences, structures and signatures of antimicrobial peptides. Nucleic acids research. 2016;44(D1):D1094–D1097.

[21] Minkiewicz P, Iwaniak A, Darewicz M. BIOPEP-UWM database of bioactive peptides: Current opportunities. International journal of molecular sciences. 2019;20(23):5978.

[22] Sanders WS, Johnston CI, Bridges SM, Burgess SC, Willeford KO. Prediction of cell penetrating peptides by support vector machines. PLoS computational biology. 2011;7(7):e1002101.

[23] Li W, Godzik A. Cd-hit: a fast program for clustering and comparing large sets of protein or nucleotide sequences. Bioinformatics. 2006;22(13):1658–1659.

[24] Gautam A, Singh H, Tyagi A, Chaudhary K, Kumar R, Kapoor P, et al. CPPsite: a curated database of cell penetrating peptides. Database. 2012;2012:bas015.

[25] Bairoch A, Apweiler R. The SWISS-PROT protein sequence data bank and its supplement TrEMBL in 1999. Nucleic acids research. 1999;27(1):49–54.

[26] Kurata H, Tsukiyama S, Manavalan B. iACVP: markedly enhanced identification of anti-coronavirus peptides using a dataset-specific word2vec model. Briefings in bioinformatics. 2022;23(4):bbac265.

[27] Řehůřek R, Sojka P. Software Framework for Topic Modelling with Large Corpora. In: Proceedings of the LREC 2010 Workshop on New Challenges for NLP Frameworks. Valletta, Malta: ELRA; 2010. p. 45–50. http://is.muni.cz/publication/884893/en.

[28] Elnaggar A, Heinzinger M, Dallago C, Rehawi G, Wang Y, Jones L, et al. Prottrans: Toward understanding the language of life through self-supervised learning. IEEE transactions on pattern analysis and machine intelligence. 2021;44(10):7112–7127.

[29] Consortium U. UniProt: a worldwide hub of protein knowledge. Nucleic acids research. 2019;47(D1):D506–D515.

[30] Hartebrodt A, Röttger R. Federated horizontally partitioned principal component analysis for biomedical applications. Bioinformatics Advances. 2022;2(1):vbac026.

[31] Wolf T, Debut L, Sanh V, Chaumond J, Delangue C, Moi A, et al. Huggingface’s transformers: State-of-the-art natural language processing. arXiv. arXiv preprint arXiv:191003771. 2019;.

[32] Berman HM, Westbrook J, Feng Z, Gilliland G, Bhat TN, Weissig H, et al. The protein data bank. Nucleic acids research. 2000;28(1):235–242.

[33] Rives A, Meier J, Sercu T, Goyal S, Lin Z, Liu J, et al. Biological structure and function emerge from scaling unsupervised learning to 250 million protein sequences. Proceedings of the National Academy of Sciences. 2021;118(15):e2016239118.

[34] Paszke A, Gross S, Massa F, Lerer A, Bradbury J, Chanan G, et al. Pytorch: An imperative style, high-performance deep learning library. Advances in neural information processing systems. 2019;32.

[35] Pedregosa F, Varoquaux G, Gramfort A, Michel V, Thirion B, Grisel O, et al. Scikit-learn: Machine Learning in Python. Journal of Machine Learning Research. 2011;12:2825–2830.

[36] McInnes L, Healy J, Melville J. Umap: Uniform manifold approximation and projection for dimension reduction. arXiv preprint arXiv:180203426. 2018;.

[37] S KPFR. LIII. On lines and planes of closest fit to systems of points in space. The London, Edinburgh, and Dublin Philosophical Magazine and Journal of Science. 1901;2(11):559–572. 10.1080/14786440109462720.

[38] Yin H, Lu Q, Wood M. Effective exon skipping and restoration of dystrophin expression by peptide nucleic acid antisense oligonucleotides in mdx mice. Molecular Therapy. 2008;16(1):38–45.

[39] Yin H, Moulton HM, Seow Y, Boyd C, Boutilier J, Iverson P, et al. Cell-penetrating peptide-conjugated antisense oligonucleotides restore systemic muscle and cardiac dystrophin expression and function. Human molecular genetics. 2008;17(24):3909–3918.

[40] Jearawiriyapaisarn N, Moulton HM, Buckley B, Roberts J, Sazani P, Fucharoen S, et al. Sustained dystrophin expression induced by peptide-conjugated morpholino oligomers in the muscles of mdx mice. Molecular Therapy. 2008;16(9):1624–1629.

[41] Betts C, Saleh AF, Arzumanov AA, Hammond SM, Godfrey C, Coursindel T, et al. Pip6-PMO, a new generation of peptide-oligonucleotide conjugates with improved cardiac exon skipping activity for DMD treatment. Molecular Therapy-Nucleic Acids. 2012;1.

